# Dissociable contributions of cortical thickness and surface area to cognitive ageing: evidence from multiple longitudinal cohorts

**DOI:** 10.64898/2026.03.20.713139

**Authors:** Ina Demetriou, Marta Correia, Didac Vidal-Piñeiro, Simon E. Fisher, Else Eising, Valentina Escott-Price, Matthew Bracher-Smith, Dace Apšvalka, Adam Attaheri, Tina Emery, Richard N. Henson

## Abstract

Cortical volume, a widely-used marker of brain ageing, is the product of two genetically and developmentally dissociable morphometric features: thickness and area. However, it remains unclear whether these two features have dissociable consequences for cognitive ageing. To address this, we analyse cross-sectional and longitudinal neuroimaging and cognitive data from one discovery cohort (Cam-CAN) and two independent, pre-registered replication cohorts (OASIS-3 and HABS-HD), leveraging wide age ranges across adulthood, different follow-up intervals and diverse population backgrounds. We show that thickness declines more steeply with age than does area, and shows stronger associations with longitudinal change in fluid cognitive abilities, fairly uniformly across the cortex. Cognitive change is also dependent on baseline thickness, independent of thickness change and independent of baseline cognitive ability. In contrast, area is comparatively stable across adulthood, at least until old age, and shows weaker and more heterogeneous associations with cognitive change. Together, these findings help to reconcile inconsistencies in the literature, and indicate that thickness provides a more sensitive marker of dynamic neurobiological processes underlying cognitive ageing, whereas area seems to reflect primarily stable, trait-like variation in cognitive ability.

## Introduction

A fundamental question in cognitive ageing research is why some individuals maintain cognitive function into late life while others show marked decline. One common candidate is cortical grey-matter volume, which shows consistent associations with cognitive decline (Oschwald et al. 2019). Yet cortical volume is a composite measure that conflates two morphometric features: thickness and area. Although both cortical features are highly heritable, they are genetically independent (Panizzon et al. 2009), follow divergent developmental trajectories (Di Biase et al. 2023), and are shaped by different neurobiological mechanisms (Rakic 2009). Understanding the relative contributions of thickness and area to cognitive decline is therefore important for developing accurate predictive models, establishing normative ageing benchmarks, and improving early detection of cognitive vulnerability in older adults. Indeed, if one cortical feature is more relevant than the other, or they make independent contributions to cognition, then relying on volume alone may obscure important patterns.

While it is widely appreciated that thickness and area should be treated separately (e.g., Bethlehem et al. 2022), the literature is surprisingly mixed on which feature is most strongly associated with cognitive decline in late life. Most studies relating brain morphometry to cognitive scores have 1) focused on the aggregate volume measure, or thickness alone, and/or 2) used cross-sectional associations across individuals, rather than longitudinal associations within individuals (see Oschwald et al. 2019 for a detailed review). Associations between longitudinal changes in brain properties and longitudinal changes in cognitive abilities offer clearer interpretation, given that they are not confounded by other individual differences (Lindenberger et al. 2011). The few longitudinal studies examining both cortical features in adulthood have reported inconsistent age-related trajectories and associations with cognitive function: Some report area to decline more with age than thickness but thickness to be the primary correlate of cognitive decline (Sele et al. 2021). Others identify thickness as a driver of age-related volumetric loss (Storsve et al. 2014), but area as the more sensitive marker of cognitive decline, with thickness relevant only in late life (Nyberg et al. 2023). These discrepancies across studies potentially reflect differences in analytic strategies, the use of regional versus global measures, differences in age ranges, or other sample characteristics. To date, no study has jointly modelled thickness and area within a single framework that estimates both cross-sectional and change-related associations, while formally testing bidirectional effects between cortical structure and cognition across the adult lifespan.

More specifically, we adopt an analytic approach that treats thickness and area as distinct routes by which exogenous factors influence cognition. Regression and mediation models inevitably encode causal assumptions, whether acknowledged or not (Rohrer 2018; Chen et al. 2024). We make these causal assumptions explicit: if area reflects stable developmental architecture shaped by early genetic profile, while thickness captures ongoing degenerative processes, then polygenic scores for intelligence (Savage et al. 2018) would be expected to influence cognition primarily through area, whereas age would operate primarily through thickness. We formalise this prediction within parallel mediation models that simultaneously estimate the independent pathways from age and polygenic scores to cognition via thickness and area. We acknowledge that stated causal assumptions cannot be directly tested within the observational design; accordingly, our analyses estimate statistical associations that would approximate causal effects only if the specified causal structure is correct and there is no unmeasured confounding.

We further use time-lagged models of longitudinal data (specifically, mixed effects models and bivariate latent change score models) to test if the data remain consistent with the assumed causal structure. By incorporating temporal precedence (Granger causality), such models provide stronger basis for inferences about the plausibility of assumed causal claims. One can ask whether baseline brain properties predict future changes in cognitive abilities, or vice versa. For example, Walhovd et al. (2022) showed that higher levels of cognitive ability were associated with reduced rates of decline in thickness, which may reflect, for example, healthier lifestyle choices that maintain brain structure. While our primary assumption is that changes in cortical features cause changes in cognitive abilities, by using bivariate latent change score models, we can simultaneously test for baseline-slope dependencies from both brain to cognition, and from cognition to brain. Again, such models do not imply causality, but can provide stronger tests of causal hypotheses (Rohrer 2018).

## Materials and Methods

### Participants

We analysed three independent adult ageing cohorts to enable discovery and replication across different sampling frameworks, age distributions, and longitudinal recruitment strategies: 1) the Cambridge Centre for Ageing and Neuroscience (Cam-CAN) cohort, with population-based lifespan sample (Shafto et al. 2014; Demetriou et al. 2025), 2) the Open Access Series of Imaging Studies III (OASIS-3) cohort, with dense multiple wave structure (LaMontagne et al. 2019), and 3) the Health and Aging Brain Study – Health Disparities (HABS-HD) cohort, with a large, community-based and ethnically-diverse sample (Petersen et al. 2025).

Cam-CAN was used as an exploratory dataset, while the subsequent analyses of OASIS-3 and HABS-HD were pre-registered here: https://osf.io/uqayb/overview (OASIS-3) and https://osf.io/mhfpk/overview (HABS-HD).

Table 1 summarises the sample characteristics, longitudinal design, cognitive measures, MRI acquisition parameters and harmonisation procedures across the three cohorts. For each cohort, we report cross-sectional and longitudinal sample sizes (before and after multivariate outlier exclusion), the number of waves available for estimating change in cognition and cortical structure, time lags between assessments, and the subsamples used for cortex-cognition analyses that require time alignment. Differences in scanners, FreeSurfer versions, and harmonisation approaches are also detailed to facilitate cross-cohort comparison.

**Table 1.** Cohort characteristics and longitudinal design across Cam-CAN, OASIS-3, and HABS-HD. Abbreviations: CU = cognitively unimpaired, PGS = polygenic score; LD = linkage disequilibrium; PCs = principal components.

|  | Cam-CAN | OASIS-3 | HABS-HD |
| --- | --- | --- | --- |
| **Cross-sectional sample** | N = 635 (319F/316M)<br>Age: 18-95<br>Without outliers: 621 | N = 1,007 (439 F/568 M)<br>Age: 42-95<br>Without outliers: 981 | N = 2,814 (1,851 F/963 M) (CU)<br>Age: 49-92<br>Ethnicity: Black (685), Hispanic (1055), White (1074)<br>Without outliers: 2,749 |
| **Timepoints used to derive change estimate** | Cognition: N = 148 with 2 waves (~12y); 105 of these with 3 waves (~6y intermediate timepoint).<br>MRI: N = 138 participants with 2 waves (~12y); 68 of these with 3 waves (~1y intermediate timepoint). | Cognition: N = 1063<br>Follow-up: 2-30 years (2-5y: 35%, 5-10y: 31%, 10-20y: 31%, >20y: 2%)<br>Waves: 2-3: 23%, 4-7: 40%, 8+: 37%<br>MRI: N = 540<br>Follow-up (2-16y): 2-5y 42%, 5-10y 44%, 10-16y 14%<br>Waves: 2-3: ~62%, 4-7: 36%, 8+: 2% | Cognition: N = 1315 (CU)<br>MRI: matched<br>Follow-up (2-6y):<br>2 waves: 65% (~2 years)<br>3 waves: 25% (~4 years)<br>4 waves: 10% (~6 years) |
| **Longitudinal sample with cognitive and cortex change estimate** | N = 138<br>Without outliers: 131 | N = 533<br>Without outliers: 511 | N = 1315<br>Without outliers: 1252 |
| **Temporally matched sample** | N = 131 (without outliers)<br>(baseline, T2: ~12 y) | N = 137 (without outliers)<br>(baseline, T2: ~6years)<br>(baseline, T2: ~3years, 50% missing, T3: ~6 years) | N = 553 (without outliers)<br>(baseline, T2: ~4 years) due to drop out T3 could not be used |
| **Cognitive measure** | Cattell Culture Fair Test | First PC of battery | First PC of battery |
| **Polygenic score (PGS)** | Fluid intelligence PGS (Savage et al. 2018); Bayesian continuous shrinkage (accounts for LD) (Ge et al. 2019); adjusted for 10 genetic PCs. | Not available | Not available |
| **FreeSurfer version** | v7.4.0 | v5.0/5.1/5.3 | v7.3.2 |
| **Scanners and acquisition sequence** | Siemens Tim Trio 3T<br>Siemens Prisma 3T<br>MPRAGE: TR=2250ms; TE=3.02ms; TI=900ms; FA=9°; voxel=1mm<sup>3</sup>; GRAPPA=2; TA=4m32s. | Siemens TIM Trio 3T<br>Sonata 1.5T / Vision 1.5T<br>MPRAGE details unavailable | Siemens Skyra 3T<br>Magnetom Vida 3T<br>MPRAGE: TR=2300ms; TE=2.98ms; TI=900ms; FA=9°; voxel=1mm<sup>3</sup>; GRAPPA=2; TA=5m12s. |
| **Harmonisation** | Not required | 1.5T vs 3T previously corrected (LaMontagne et al. 2019) | Longitudinal ComBat*<br>(Harmonisation model: baseline age, time lag, 2-way interaction; covariates: ethnicity, sex, education). |

\* (Beer et al. 2020; https://github.com/jcbeer/longCombat)

### Cognitive measures

We focused on fluid intelligence, often indexed by the first principal component of scores across a range of standardised cognitive tests (Spearman’s ‘*g*’; Spearman 1961). This was the approach taken in the OASIS-3 and HABS-HD cohorts (details below), though for Cam-CAN, we used the sum score from the Cattell Culture Fair test, an abstract reasoning test that normally loads highly on the first principal component (since Cam-CAN did not have a comparable battery of standard tests). Therefore, we assume the Cattell test fairly approximates *g* and use the terms ‘g’ and ‘cognition’ interchangeably.

For OASIS-3, the cognitive tests were two Category Fluency tests (Animals and Vegetables; Acevedo et al. 2000), the Digit Symbol Substitution Test (DSST; Wechsler 1955), two Trail Making Tests (TMT A & B; Reitan & Wolfson 1995) and the WAIS Block Design test (Wechsler, 1981). For TMT A and TMT B, we removed values equal to or above 150 sec and 180 sec respectively as a discontinuation time. For HABS-HD, the cognitive tests were a Phonemic Fluency Test (F-A-S, Spreen 1977), the Animal Fluency Test, the Digit Symbol Substitution Test, two Trail Making Tests and the Digit Span Test (Wechsler 1981). We used all available cognitive tests that traditionally load on *g* and were missing less than 30% measures across all waves (as pre-registered). Missing test scores (<30%) were imputed with fully conditional specification (FCS); specifically, using 2-level predictive mean matching (PMM), implemented by 2l.pmm method in R from the *“miceadds”* package, a robust technique for unbiased imputations in longitudinal data with irregularly spaced assessments (Wijesuriya et al. 2025).

Cognitive tests were individually adjusted for practice effects and, in Cam-CAN, additionally for format differences. Adjusted scores were derived as residuals of a mixed-effects model including fixed effects of practice and format and a participant random intercept. Practice was modelled as asymptotic after the first exposure (Staff et al. 2014). For OASIS-3 and HABS-HD, we assumed measurement invariance, and derived a weighted composite score (*g*) by multiplying the matrix of adjusted cognitive scores (across all waves) by the extracted PC1 loading vector.

To address the possibility that age-related acceleration means that baseline age confounds estimates of change slopes, we additionally derived cognitive scores adjusted for nonlinear age effects. Following Capogna et al. (2025), the adjusted scores were the residuals of a Generalised Additive Mixed Model (GAMM) that used splines to capture effects of age and a participant random intercept (plus format and practice as fixed effects). However, because age is intrinsically linked to the biological processes underlying both cortical and cognitive decline, such adjustment may also remove meaningful age-dependent biological signal, particularly in lifespan cohorts where age-related variance constitutes a substantial component of true inter-individual differences in change. We therefore report age-adjusted analyses as sensitivity checks in the Supplementary Materials, while presenting unadjusted models as primary results.

### MRI measures

We used global mean cortical thickness (mm) (thickness) and total cortical area (mm²) (area) from FreeSurfer processing of the T1-weighted MRI images. When estimating cortex changes, we repeated analyses after adjustment for nonlinear age effects, analogous to cognitive changes above, and as reported in Supplementary Materials.

### Genotyping data

DNA samples were genotyped using the Illumina Infinium “OmniExpressExome” array, filtered for minor allele frequency < 0.05, missingness < 0.05, Hardy Weinberg p < 1×10^−6^ and non-European ancestry according to principal component analysis with 1000 genomes data, and imputed using the Haplotype Reference Consortium version 1.1 panel (Henson et al. 2020).

### Polygenic scores for fluid intelligence

A polygenic score (PGS) for fluid intelligence was based on pooling effect sizes across many single nucleotide polymorphisms (SNPs) identified in a GWAS study of other independent cohorts (Savage et al. 2018). It was computed using the Bayesian continuous shrinkage prior approach with PRS-CS (1KG LD reference), which robustly infers the posterior effect sizes of SNPs, while also taking into account the external linkage disequilibrium (Ge et al. 2019). Individual-level PGSs were then calculated for all Cam-CAN participants, adjusted for the first 10 genetic principal components (to control for population stratification effects), and then standardised.

### Statistical analyses

#### Outliers

For all analyses, multivariate outliers were identified using Mahalanobis distance (Leys et al. 2018). Observations exceeding the 97.5th percentile of the χ² distribution were excluded. Cross-sectional models included baseline cognition, thickness, area, age, sex, years of education, and PGS (where available). Longitudinal models included change of cognition, thickness, and area, along with baseline age, sex, and years of education.

#### Cross-sectional parallel mediation (SEM)

To test whether thickness and area represent dissociable pathways from age and genetics to cognition, we formalised our causal hypotheses using path analysis, illustrated in Figure 1. Age and PGS (Cam-CAN only) were treated as exogenous variables. Thickness and area were specified as parallel mediators, and cognition as the distal outcome. Note that, while thickness is the average over the cortex, area is the total across the cortex, and is therefore related to total brain size. In case any mediation effects owed to differences in cortical area/thickness/fluid intelligence between men and women, we included sex as a predictor of all three. However, given uncertainty about whether more years of education (YoE) causes higher cognitive scores, or whether genetics or early-life developmental factors determine cognitive potential that then causes more years in education (or both; Fjell et al. 2025), we tested models with and without YoE. More specifically, we added a path to YoE from area (as proxy for brain size), to capture possibility that YoE is determined by genetic/early-life factors, but also a direct path from YoE to *g*, to capture the possibility that education contributes to cognition via routes other than brain size (analogous to sex variable above). The results with YoE are reported in Supplementary Table 1.

**Figure 1.**
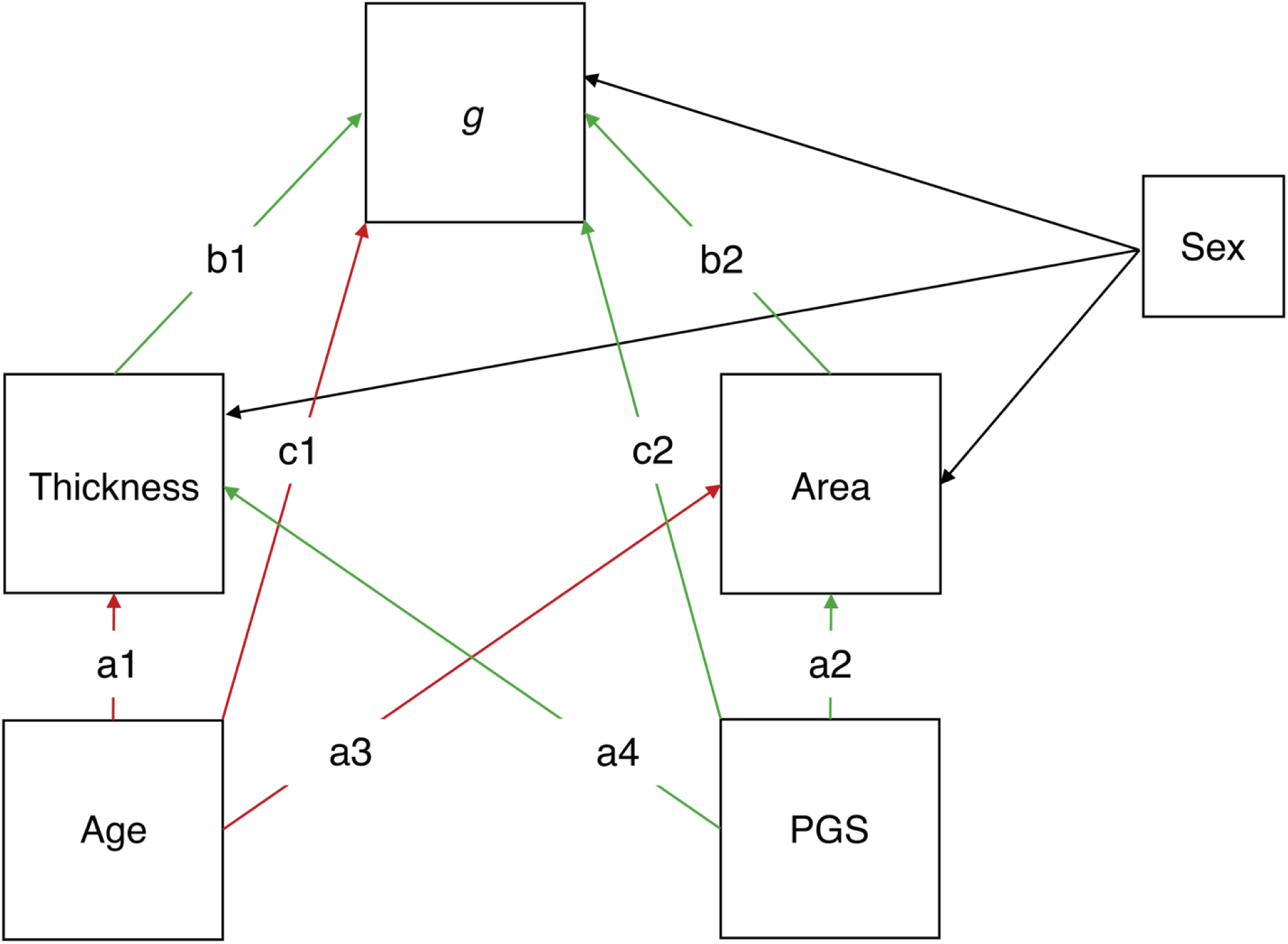
Directed acyclic graph illustrating a structural equation model (SEM) testing dissociable pathways from age and polygenic score (PGS) to g via thickness and area. Colour denotes expected positive (green) or negative (red) relationship between the variables of interest; black – to/from confounding variables.

We did not include intracranial volume (ICV) as a predictor of g because we assumed that any effect of head size must be mediated by cortical area or thickness (see Chen et al. 2024, for further discussion of such causal assumptions). However, we did fit another model in which ICV was a predictor of area and thickness (i.e., effectively adjusting these GM measures by ICV; Fürtjes et al. 2024). This affects the “a” paths in Figure 1, and hence the mediation effect sizes (which are the product of “a” and “b” paths). The results of this model are shown in Supplementary Table 2.

However, it is possible that including ICV as a predictor of area and thickness could attenuate true mediation effects. For example, it is possible that more overall GM area does cause better cognition, and therefore adjusting for ICV will reduce the true mediation by area (or thickness), by virtue of reducing the “a” paths (e.g., from PGS). Indeed, ICV was positively correlated with g in two of the cohorts (CamCAN: r = 0.28, p <.001, OASIS-3: r = −0.06, p = 0.07, HABS-HD: r = 0.20, p<.001). Secondly, ICV was also correlated with age, with varying magnitudes and directions across cohorts (CamCAN: r = −0.16, p < .001; OASIS-3: r = 0.11, p < .001; HABS-HD: r = 0.12, p < .001), which could affect the “a” paths from Age. For these reasons, the main Results come from models without ICV.

We defined direct paths from age to thickness (a1), area (a3), and cognition (c1), and from PGS to thickness (a4), area (a2), and cognition (c2). Paths from thickness and area to cognition were defined as b1 and b2, respectively. Indirect effects were calculated as products of path coefficients (e.g., a1 × b1, a2 × b2, a3 × b2 and a4 × b1). The proportion of variance mediated was computed relative to the total effect. To formally test dissociation between thickness and area pathways, we specified contrast variables as the difference between the a1 × b1 and a3 × b2, and the difference between a2 × b2 and a4 × b1 for PGS, and estimated the standard error for these contrasts through 5000 bootstrap resamples.

#### Longitudinal change in cortex structure

To estimate age-related change in thickness and area, we fitted linear mixed-effects models: *GM measure ∼ A0 + dA + A0:dA + (1|Participant)*, where A0 denotes baseline age and dA denotes time since baseline. This age decomposition is helpful to separate between-participant differences, which could be susceptible to cohort effects and selective survival bias (Raz and Lindenberger 2011), from true within-participant longitudinal change. The interaction term tests whether rate of change depends on baseline age (i.e., accelerated decline). Raw units were retained (mm for thickness; mm² for area) to characterise biologically interpretable loss of cortical measures per year. Standardised coefficients are additionally reported to compare relative effect sizes between thickness and area. In HABS-HD, wave structure allowed, we additionally conducted pattern-mixture analysis, extending the mixed-effects models to include attrition pattern group and its interactions with time (e.g., Pattern Group:dA), where Pattern Group was defined based on the last available timepoint. This approach allowed us to test if longitudinal estimates were influenced by differential dropout.

#### Estimation of individual slopes

We estimated the individual change (slope) for each participant by the linear coefficient of the simple regression of *g* / thickness / area against the lag between the 2-3 timepoints for that participant. Note that the scores on each cognitive task were first adjusted for practice / format effects, using the participant-wide mixed effect models described above (while the analyses in Supplementary Tables 3 and 4 also adjusted *g*, thickness and area for nonlinear effects of age).

To test if the rate of cognitive decline was associated with the rate of thickness or area decline, we used a simple regression between the change estimates. We used Steiger’s test to formally test if these relationships differed significantly in their strength. To test whether the relationship with cognitive change was different for thickness and area, and also whether the cognitive change depended on past (baseline) grey-matter values, we fit an extended model:

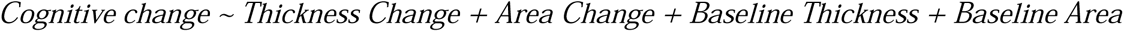

We used a linear hypothesis test to formally compare the relative size of the main effects of thickness change and of area change.

#### Bivariate Latent Change Score Models (BLCSMs) & Latent Growth Models (LGMs)

Better evidence for temporal relationships between baseline values and slopes can be tested with a bivariate latent change score model (BLCSM) (Kievit et al. 2018). This model extends the above mixed effects model by also allowing for lagged relationships from baseline cognition to changes in thickness or area. While one might expect past cortex state to predict future cognitive ability, the reverse is also possible, i.e., that people with higher baseline cognitive ability might show less cortex ageing owing, for example, to healthier lifestyle choices, which in turn can lead to accumulation of unmeasured brain and cognitive changes before the first measurement occasion. BLCSMs can model both lagged effects simultaneously.

Unlike the above mixed effects models however, BLCSMs assume a fixed lag for all participants and a fixed lag across all timepoints. The latter necessitated using data from only two timepoints, namely baseline and the latest (or in case of OASIS and HABS-HD, the most available) follow-up (see Table 2). We used raw values, not adjusted for effects of practice or format, because BLCSM models rely on between-person variance and therefore removing participant-specific random effects would make change-change associations and baseline-change effects uninterpretable. However, to adjust for potential nonlinear effects of age, Supplementary Table 5 also reports models with baseline age as a predictor of latent change in cognition and latent change in cortex, comparable to the GAMM residualisation described above for mixed effects models.

In OASIS-3 and HABS-HD, which had sufficient numbers of participants with 3 timepoints, we also fit Latent Growth Models (LGMs), since they provide more robust estimates of linear slopes than BLCSMs (Kievit et al. 2018). The convergence between these 3-wave LGM results and the 2-wave BLCSM results are shown in Supplementary Table 6 and Supplementary Figure 1.

Table 2 overviews the main research questions addressed, along with analyses type and key variables.

**Table 2.**
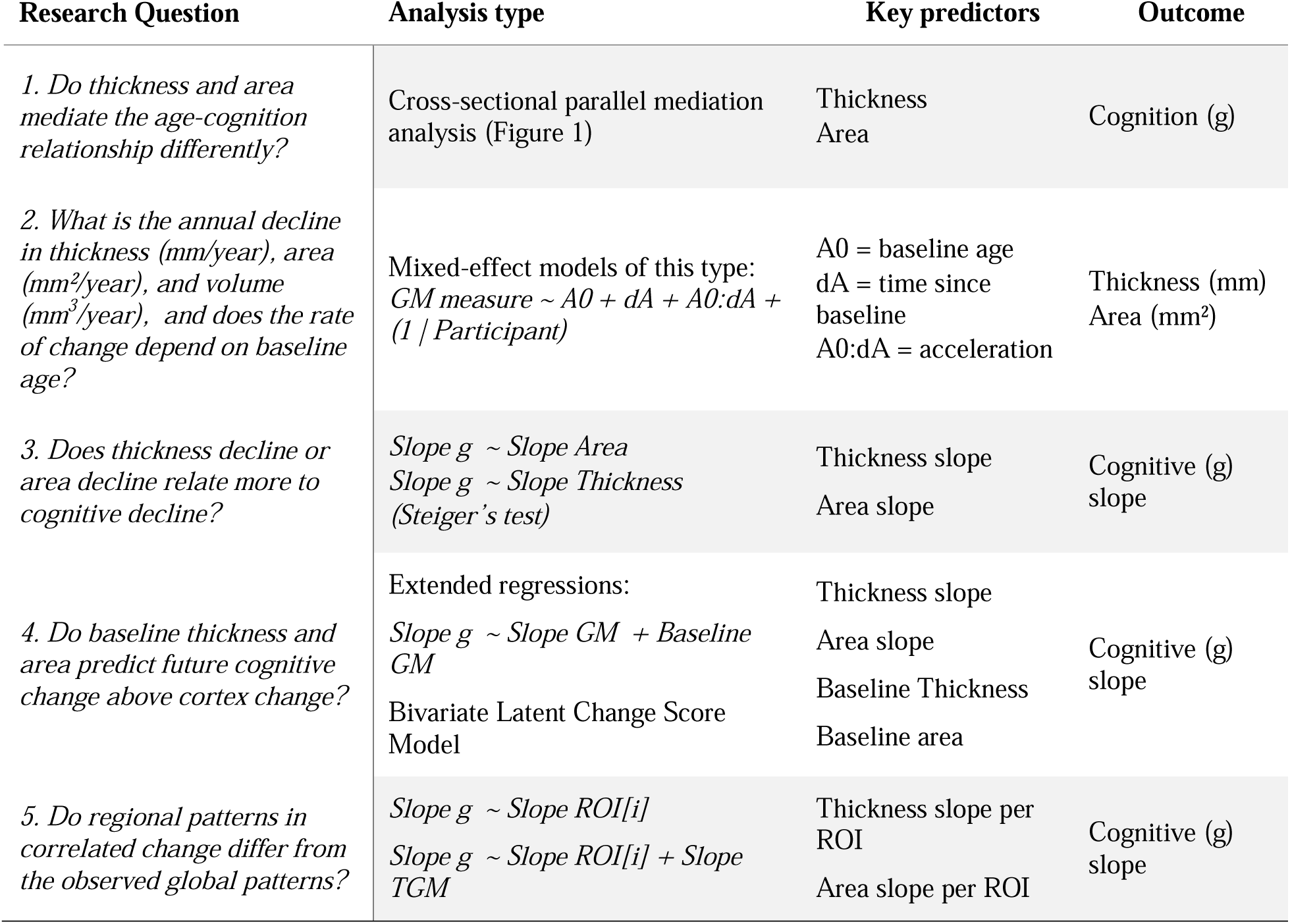
Summary of the main research questions with analysis type used to answer it, key predictors and outcome variable for each. The Results section follows these questions sequentially.

### Deviations from Pre-registration

The pre-registered analyses for OASIS-3 and HABS-HD were followed with the following deviations:

1. **Slope estimation.** Individual slopes were estimated using simple per-participant regression rather than mixed-effects random slopes (ranef). The reason is that the group-regularisation inherent in the random slopes will differentially affect individuals with 2 versus 3 timepoints (and in fact bias estimates for individuals with any other under-represented demographic characteristic); deriving slopes individually, albeit noisier, ensures comparability of change estimates across participants. Nonetheless, results using the pre-registered approach are available online and did not differ from those reported.
2. **LGM specification.** The pre-registered three-wave LGMs were only fitted as supplementary models. Two-wave BLCSM were used as the primary analysis instead for direct comparability with the Cam-CAN cohort. LGM models reported in the Supplementary Materials showed the same pattern of effects.
3. **Cognitive score derivation.** The pre-registered CFA/FIML approach was replaced by a weighted PC1 composite, with missing scores imputed via two-level predictive mean matching, more appropriate for irregularly spaced longitudinal data.
4. **Practice effect adjustment.** Cognitive scores were adjusted for practice effects prior to PCA; this was not mentioned in the pre-registration.
5. **Extended regression model.** A regression of the form cognitive change ∼ cortical change + baseline cortex was added to test whether baseline thickness predicted future cognitive change above concurrent change; this was not pre-registered but is consistent with the BLCSM and LGM findings.

## Results

### Cross-sectional parallel mediation

Both thickness and area significantly and independently mediated the relationship between age and cognition, but thickness was a consistently stronger mediator (Table 3). In Cam-CAN, thickness mediated 14% of the total age effect on cognition, while area mediated only 9%. This pattern replicated in OASIS-3 (29% vs 13%) and HABS-HD (19% vs 15%). The contrast between the proportion of variance in g mediated by thickness versus area was significant in OASIS-3 (contrast = 0.16, p <.001), though did not reach significance in Cam-CAN (contrast = 0.05, p = .213) or HABS-HD (contrast = 0.05, p = .229).

**Table 3.** Indirect effects of age (and polygenic score) on cognition through thickness and area across cohorts.

| Cohort | Mediator | Estimate (% mediated), p-value |  |
| --- | --- | --- | --- |
|  |  | Age → g | PGS → g |
| Cam-CAN<br>(N = 621) | Thickness | $\beta=0.10$ (**14%**), $p<.001$ | $\beta=0.008$ (5%), $p=.086$ |
| | Area | $\beta=0.06$ (**9%**), $p<.001$ | $\beta=0.021$ (**12%**), $p=.005$ |
| OASIS-3<br>(N = 981) | Thickness | $\beta=0.15$ (**29%**), $p<.001$ | NA |
| | Area | $\beta=0.07$ (**13%**), $p<.001$ | |
| HABS-HD<br>(N = 2749) | Thickness | $\beta=0.04$ (**19%**), $p<.001$ | NA |
| | Area | $\beta=0.03$ (**15%**), $p<.001$ | |

The genetic dissociation showed the reverse structural pattern (Table 3): Area mediated 12% of the PGS effect on cognition, while thickness showed no significant mediation. However, given that the direct contrast between these mediations did not reach significance (contrast = −0.07, p = .145), and that PGS were not available in OASIS-3 or HABS-HD, these results need to be replicated in future, larger studies.

### Longitudinal decline in cognition and cortex

Thickness and area both showed marked age-related reductions (Table 4, Figure 2), but differed in their patterns of decline. Thickness declined linearly and consistently across cohorts (−0.004 to −0.005 mm/year), while area showed greater heterogeneity in raw decline rates (ranging from approximately −190 to −564 mm²/year). Pattern-mixture analysis in HABS-HD did not show any evidence that selective attrition biased these estimates for either thickness (Pattern Group:dA, β = −0.001 mm/year, p = .222) or area (Pattern Group:dA, β = −41.13 mm^2^/year, p = .312). Volume showed robust age-related decline across cohorts, but no consistent advantage over thickness (only a slight advantage in OASIS-3). Volume effects generally fell between those for thickness and area (Cam-CAN and HABS-HD), suggesting that combining the two measures may attenuate rather than increase sensitivity to age-related cortical change.

**Table 4.** Cross-sectional (A0), linear (dA), and nonlinear (A0:dA) effects of age on thickness (mm), area (mm²) and volume (mm^3^) across cohorts (A0 = baseline age (years); dA = time since baseline at each measurement occasion (years)). Partial marginal R^2^ (in brackets) indicates the proportion of unique variance of data explained by fixed effects.

| Measure | Effect | Cam-CAN (n=131) | OASIS-3 (n=511) | HABS-HD (n=1252) |
| --- | --- | --- | --- | --- |
| **Thickness** | A0 ( mm/year) | -0.004 (37.7%), $p < .001$ | -0.005 (17.5%), $p < .001$ | -0.003 (9.7%), $p < .001$ |
| | dA (mm/year) | -0.004 (4.4%), $p < .001$ | -0.005 (2.6%), $p < .001$ | -0.005 (1.1%), $p < .001$ |
| | A0:dA (mm<sup>2</sup>/year<sup>2</sup>) | -0.00004 (<1%), $p = .070$ | -0.0003 (1.4%), $p < .001$ | -0.0001 (<1%), $p = .012$ |
| **Area** | A0 (mm<sup>2</sup>/year) | -228.10 (3.2%), $p = .036$ | -220.22 (<1%), $p = .007$ | -100.85 (<1%), $p = .063$ |
| | dA (mm<sup>2</sup>/year) | -189.18 (<1%), $p < .001$ | -563.71 (1.4%), $p < .001$ | -477.63 (<1%), $p < .001$ |
| | A0:dA (mm<sup>2</sup>/year<sup>2</sup>) | -1.97, (<1%), $p = .068$ | -15.57 (<1%), $p < .001$ | -11.08 (<1%), $p < .001$ |
| **Volume** | A0 (mm<sup>3</sup>/year) | -1466.04 (16.5%), $p < .001$ | -1419.69 (6.3%), $p < .001$ | -793.19 (2.2%), $p < .001$ |
| | dA (mm<sup>3</sup>/year) | -1026.25 (1.1%), $p < .001$ | -2136.58 (2.9%), $p < .001$ | -2117.65 (<1%), $p < .001$ |
| | A0:dA (mm<sup>3</sup>/year<sup>2</sup>) | -6.73 (<1%), $p = .117$ | -60.59 (<1%), $p < .001$ | -38.150 (<1%), $p < .001$ |

**Figure 2.**
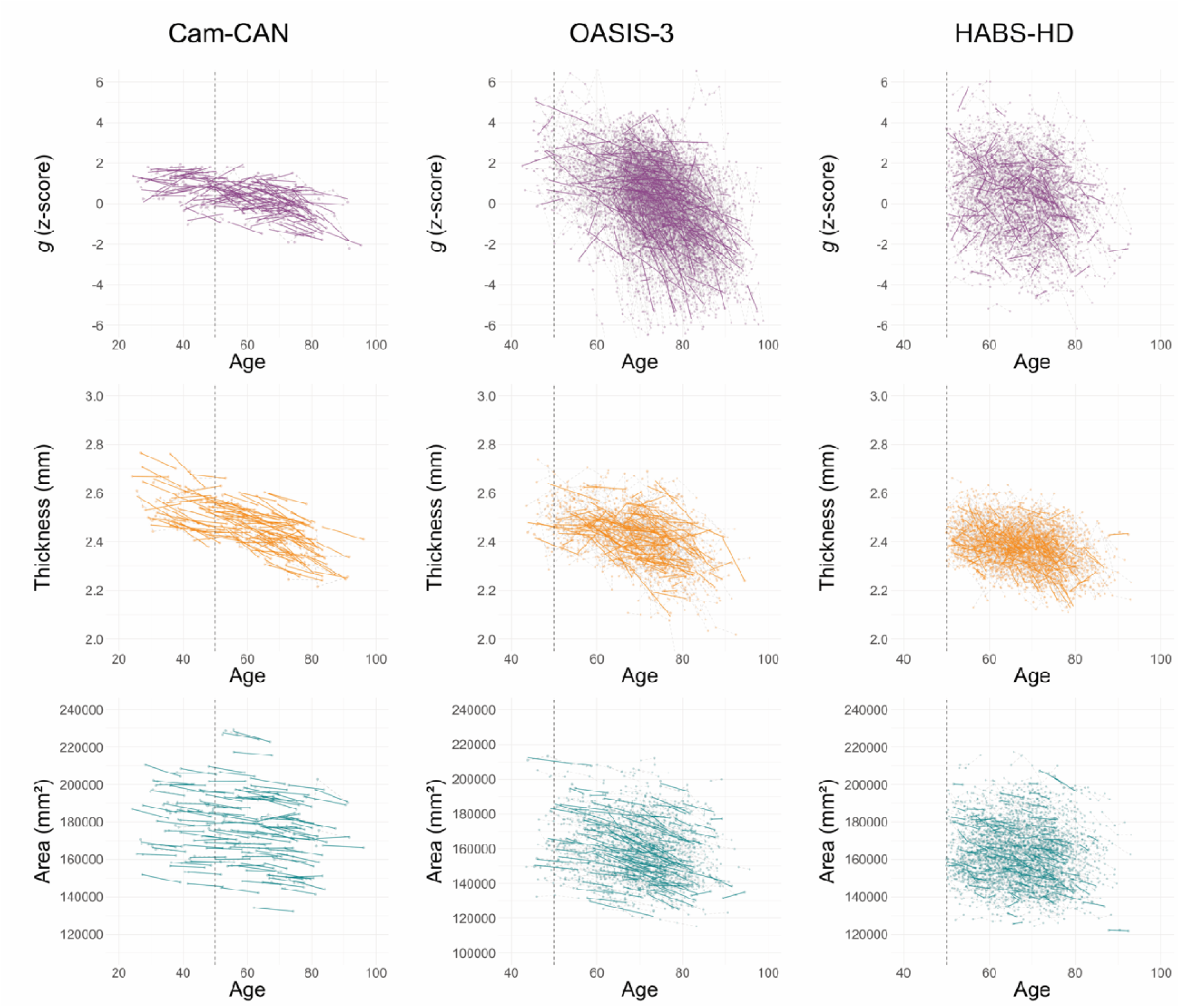
Longitudinal trajectories of g (z-score), thickness (mm) and area (mm^2^) across cohorts. The vertical dashed line marks age 50, corresponding to the approximate lower age bound of the ‘oldest’ cohort (HABS-HD), and thus a reference point for visual comparison across cohorts. The plots ‘highlight’ randomly selected n = 131 participants in OASIS-3 and HABS-HD (to match the number in Cam-CAN) for visual comparison only.

### Correlated longitudinal cortex-cognitive change

Correlated change analyses further supported a dissociation between the relationships between thickness and area on cognition (Table 5, Figure 3). Across cohorts, change in thickness was positively associated with change in cognition, accounting for ∼3% of the variance in cognitive decline (irrespective of any adjustment for nonlinear age effects; see Supplementary Table 3 and 4). In contrast, change in area was not reliably related to cognitive change and, where present, accounted for less than 1% of variance. In OASIS-3 at least, Steiger’s test confirmed that this correlation was significantly stronger for thickness than area. These results remained robust to the adjustment for nonlinear age effects (Supplementary Table 3). Volume change was also associated with cognitive change, but showed no consistent advantage over thickness. Overall, volume explained little additional variance in cognitive change (in HABS-HD only) and generally showed effect sizes between those for thickness and area (in Cam-CAN and OASIS-3), consistent with attenuation of the thickness-related cognitive signal by the inclusion of area.

**Figure 3.**
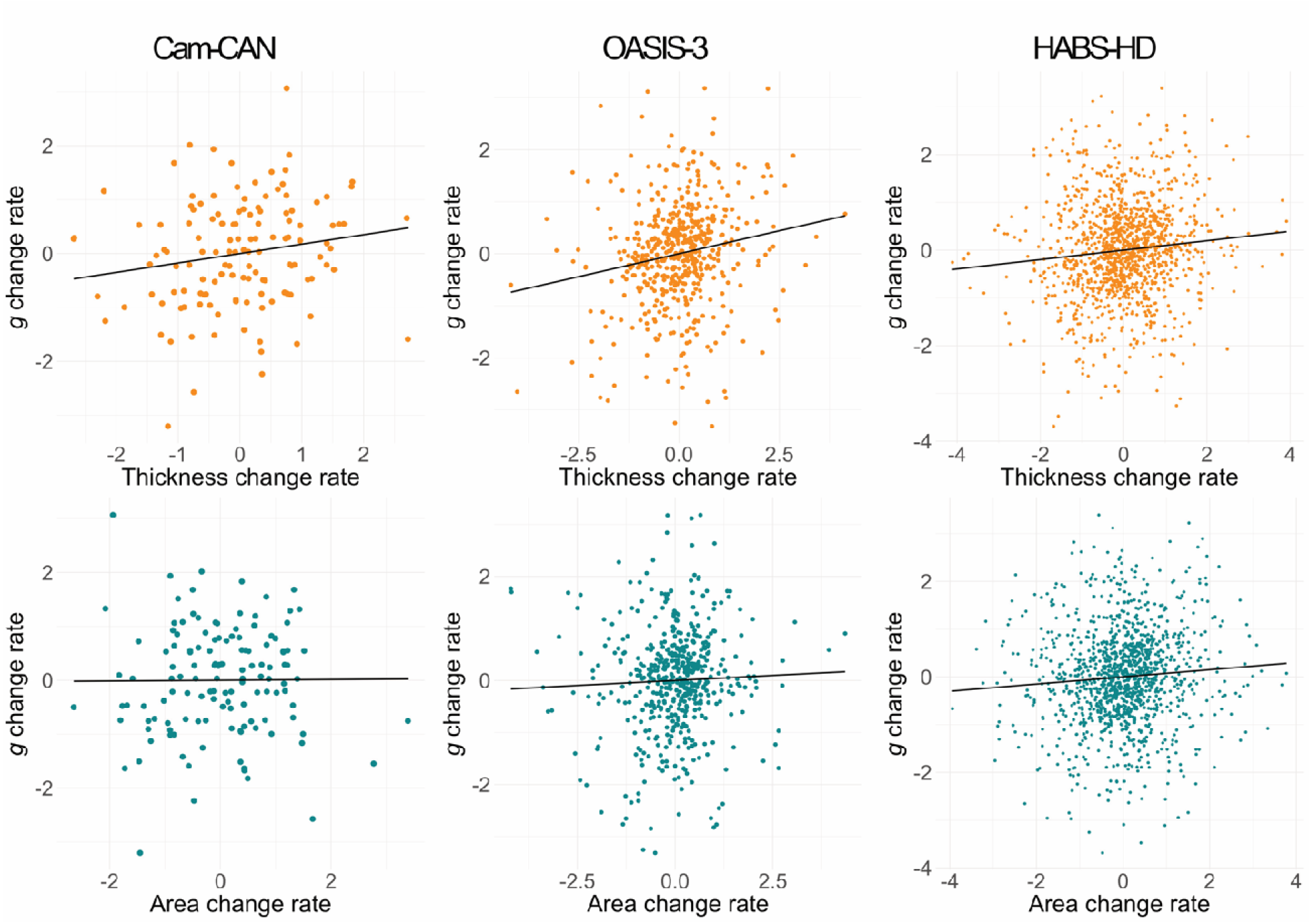
Scatterplots showing associations between thickness change rate (top row) and area change rate (bottom row) with g change rate across cohorts. All measures z-scored.

**Table 5.**
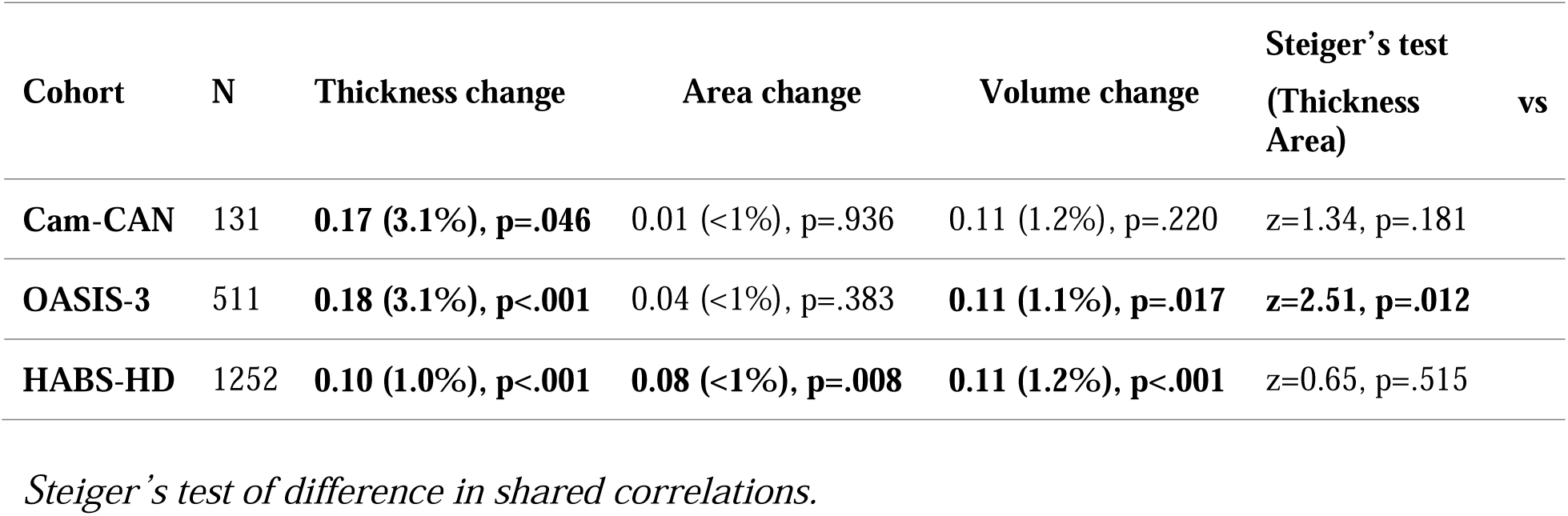
Standardised effect sizes (β) from simple regression models with one predictor (thickness change or area change). R^2^ (in brackets) indicates proportion of variance explained.

### Dependency of cognitive change on baseline cortex

Going beyond simple, independent change-change correlations, we fit a combined model that predicted cognitive change from change in both thickness and area, as well as from baseline values of thickness and area (see Methods). As found previously, thickness change, but not area change, predicted cognitive change in all cohorts, but now above any effects of baseline, and significantly more so in OASIS-3 (according to a linear hypothesis test; Table 6). More importantly, baseline thickness made an independent contribution to cognitive decline, accounting for ∼1-6% of additional variance above concurrent change effects, whereas baseline area did not contribute to cognitive change.

**Table 6.** Correlated change and baseline cortex – cognition change associations from extended longitudinal models. Linear Hypothesis Tests (LHT) indicate whether change–change associations differ significantly between thickness and area.

|  | N | Thickness<br>change | Thickness<br>baseline | Area<br>change | Area<br>baseline | LHT |
| --- | --- | --- | --- | --- | --- | --- |
| **Cam-CAN** | 131 | **0.18 (3.4%),**<br>**p=.037** | **0.24 (6%),**<br>**p=.005** | 0.004 (<1%),<br>p=.964 | 0.08 (<1%),<br>p=.335 | F=2.09,<br>p=.150 |
| **OASIS-3** | 511 | **0.18 (3.4%),**<br>**p<.001** | **0.23 (5.5%),**<br>**p<.001** | -0.01 (<1%),<br>p=.766 | -0.02 (<1%),<br>p=.576 | **F=8.33,**<br>**p=.004** |
| **HABS-HD** | 1252 | **0.12 (1.3%),**<br>**p<.001** | **0.12 (1.4%),**<br>**p<.001** | 0.05 (<1%),<br>p=.061 | -0.004 (<1%),<br>p=.889 | F=2.21,<br>p=.137 |

This pattern mostly held with the adjustment for nonlinear age effects (Supplementary Table 4): baseline thickness remained significant with the GAMM correction in OASIS-3 and HABS-HD, but was attenuated in Cam-CAN. This attenuation likely reflects the removal of nonlinear age variance that overlaps with the baseline thickness effect due to cohort’s wide age range (18-95 years), compared to the more age-restricted samples in OASIS-3 (42-95) and HABS-HD (49-92).

### Mutual dependency of change on baseline

While the previous analysis demonstrated lagged effects of baseline thickness on subsequent cognitive change, the reverse might also be possible, i.e., baseline cognitive levels could predict subsequent cortical decline (Walhovd et al. 2022), e.g., through healthier lifestyle choices. To allow for such symmetric, lagged effects, we used 2-wave BLCSMs and 3-wave LGMs (see Methods). Though these methods reduced the sample size (since only matched timepoints for cognitive and MRI assessments can be used), they replicated the main findings above, as shown in Figure 4 and Table 7 (for BLCSMs; for LGMs see Supplementary Table 6).

**Figure 4.**
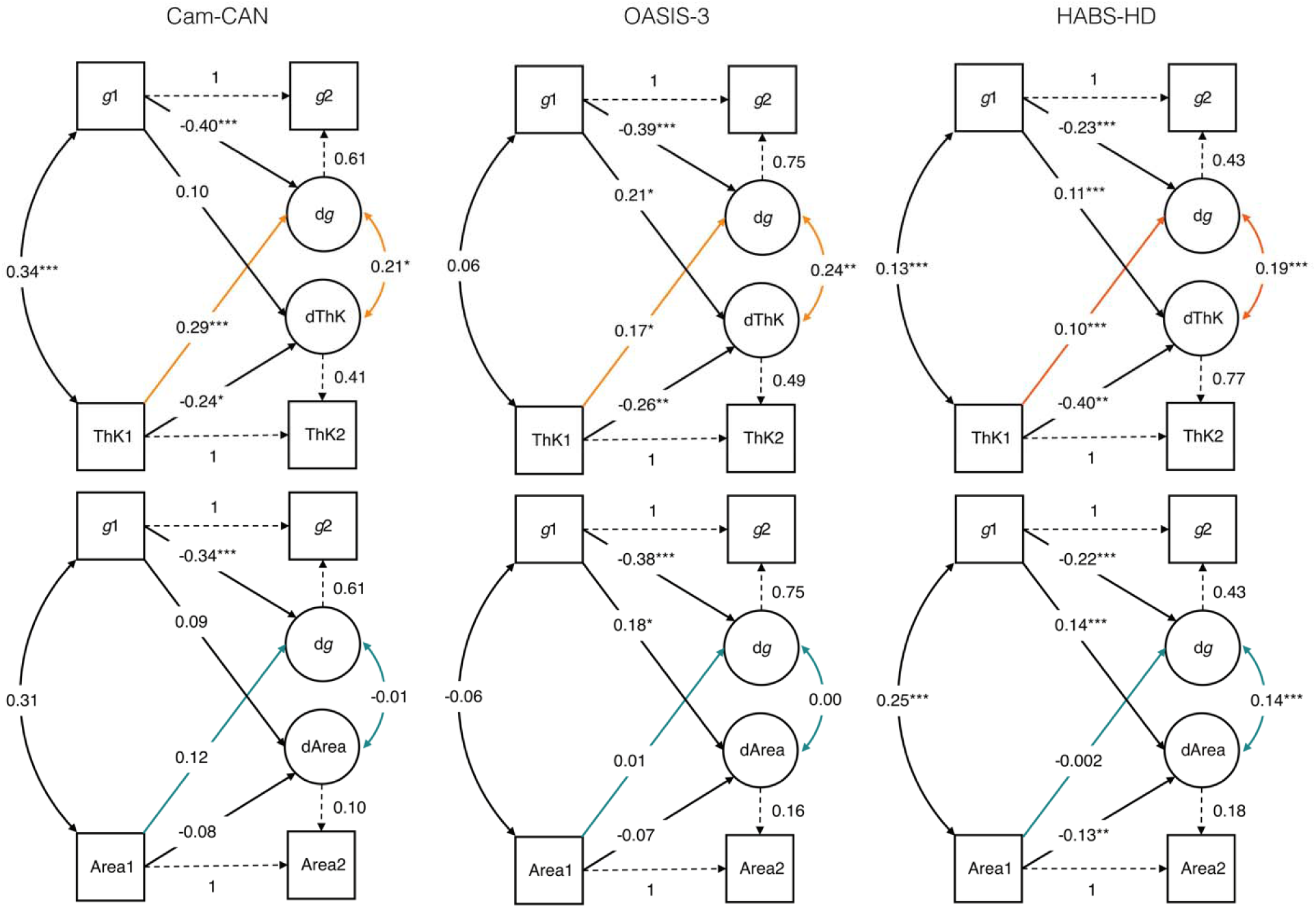
Visual representation of BLCSM results. Rectangles – observed variables at baseline (T1) and latest available follow-up (T2); circles – latent change factors (dg; dThK (thickness); dArea). Curved double-headed arrows – covariances between latent change factors. Dashed paths indicate parameters that were fixed for model identification. Coloured paths (orange: thickness models; turquoise: area models) represent the primary effects of interest, capturing cross-domain influences between baseline levels and subsequent change. Values shown are standardised parameter estimates. Statistical significance is indicated as follows: p < .05 (*), p < .01 (**), p< .001 (***).

**Table 7.** Bivariate latent change score model results of latent change covariance and cross-lagged relationships between baseline and change measures.

| Cohort | N | Measure | $\Delta \text{Cortex} - \Delta g$ | Baseline Cortex - $\Delta g$ | Baseline cog - $\Delta \text{Cortex}$ |
| --- | --- | --- | --- | --- | --- |
| **Cam-CAN** | 131 | Thickness | **0.21, p=.013** | **0.29, p<.001** | 0.10, p=.303 |
|  |  | Area | -0.01, p=.927 | 0.12, p=.136 | 0.09, p=.250 |
| **OASIS-3** | 137 | Thickness | **0.24, p=.006** | **0.17, p=.034** | **0.21, p=.010** |
|  |  | Area | 0.003, p=.965 | 0.01, p=.874 | **0.18, p=.039** |
| **HABS-HD** | 553 | Thickness | **0.19, p<.001** | **0.10, p=.027** | **0.11, p=.007** |
|  |  | Area | **0.14, p=.002** | -0.002, p=.968 | **0.14, p=.002** |

Correlated change estimates from BLCSMs were comparable to regression analyses across cohorts, despite reduced power. Likewise, baseline thickness predicted subsequent cognitive change, whereas baseline area did not, reinforcing the directional asymmetry central to the age-related dissociation between these two cortical features.

Reverse lagged effects (from baseline cognition to cortical change) were more variable across samples and model specifications. In Cam-CAN, baseline cognition did not predict subsequent change in thickness or area. In OASIS-3 and HABS-HD baseline cognition predicted change in both metrics (replicating Walhovd et al. 2022). Critically, no cohort showed that baseline area predicted cognitive change, irrespective of model specification whereas baseline thickness did so consistently.

Adjustment for baseline age reduced some cross-lagged effects in these BLCSMs (Supplementary Table 5), but did not alter the central dissociation: thickness mostly retained prospective associations with cognitive change, whereas area did not. Together, these findings strengthen the interpretation that thickness is more tightly linked to dynamic ageing-related processes, whereas the relationship between area and cognition is less linked to such dynamic state effects, and relatively more to static trait effects.

### Regional results

Given that our results from global measures of thickness and area differed from those reported by Nyberg et al. (2023) for their three ROIs (rostral middle frontal, middle temporal, inferior parietal), we repeated our analyses for these ROIs based on the Desikan-Killiany atlas. In Cam-CAN, neither thickness nor area change was significantly associated with cognitive change in the three Nyberg ROIs (see Supplementary Table 7). In OASIS-3, thickness change was significantly associated with cognitive change in middle temporal and inferior parietal, but not rostral middle frontal; area estimates were not available in OASIS-3 for ROIs. In HABS-HD, thickness change was associated with cognitive change in inferior parietal and middle temporal, but not rostral middle frontal; for area, middle temporal and rostral middle frontal showed a significant association, whereas inferior parietal did not. When examined across all 34 Desikan–Killiany regions, thickness-cognition associations were directionally consistent across ROIs (even when not always surviving correction for multiple comparisons), whereas area-cognition associations were more heterogeneous across ROIs (including changes in sign) and rarely survived correction (see Supplementary Table 7). This suggests that the cognitive consequences of declines in thickness are likely to be fairly uniform across the cortex (supporting our use of a global measure in the above analyses), whereas any consequences of declines in area (if any) are likely to be restricted to a few parts of cortex. We also report regional values controlled for global thickness and area (see Supplementary Table 8), because prior work on white matter has shown that a general factor accounts for substantial shared variance across tracts, suggesting that region-specific effects should be examined after partialling out global effects (Penke et al. 2010). The ROIs showing corrected effects changed somewhat, but almost no regions survived the FDR correction.

## Discussion

We tested a causal framework in which thickness is hypothesised to index dynamic, ageing-related processes, while area is hypothesised to reflect earlier, genetically-influenced neurodevelopmental factors, largely preserved throughout adulthood. This model assumes thickness is a mediator of the causal pathway linking ageing-related processes to cognitive decline, whereas area is a mediator on the pathway linking early genetic influences to baseline cognitive differences. Although the causal assumptions embedded in our model cannot be directly tested within observational data, the pattern of results across i) cross-sectional mediation, ii) longitudinal slope-slope, and iii) time-lagged bivariate analyses, across three independent healthy ageing cohorts, was largely consistent with this framework.

The cross-sectional, parallel mediation analyses showed that thickness is a stronger mediator of the age-cognition relationship than is area. In contrast, polygenic scores for cognition (though only available in the Cam-CAN cohort) were mediated by area, but not thickness, consistent with evidence that area is more strongly influenced by genetics and early developmental factors. Together, these findings support a dissociation between thickness and area in their associations with cognition.

Longitudinal analyses provided stronger evidence for differential role of these morphometric features. Thickness decline was robust and approximately linear across cohorts, and its rate of change was positively associated with cognitive decline, accounting for approximately 1-3% of variance. While modest, it is important to note that we only focused on cortical measures, whereas cognitive ageing is likely to be a multifactorial process, influenced by other types of brain change, such as changes in white-matter, functional connectivity, etc. For example, individuals with severely reduced cortical tissue due to hydrocephalus can nonetheless display normal cognitive function (Lewin 1980). This underscores the importance of measuring multiple brain properties in order to understand the puzzle of “cognitive reserve” (Henson 2026). Nonetheless, the consistency of thickness effects across the three cohorts suggests that these modest longitudinal associations are reliable and have implications for future research, and that analyses relying solely on volume may obscure dissociable relationships relevant for establishing individual ageing trajectories

Importantly, baseline thickness predicted future cognitive change, even when controlling for reverse paths from cognition to cortex structure in the BLCSM and LGM models, whereas no such lagged effect was observed for area in any of the cohorts or modelling techniques. These time-lagged findings are consistent with the interpretation that thickness is more closely coupled to age-related cognitive change. Importantly across all cohorts and types of analyses, the effects of thickness and area remained largely consistent even after various adjustments for nonlinear age effects in cortex and cognition. In some samples, higher baseline cognition was associated with reduced subsequent decline in thickness or area, although these effects were less consistent. Any cognition-related influence on structural change is likely to be indirect, e.g. through healthier lifestyle, or other third variable (Walhovd et al. 2022).

Note that we mostly focused on global measures, leaving open the possibility of regional or domain-specific dissociations that may differ from the global pattern (e.g. volume patterns in Cox et al. 2021). However, our analyses of regional variation (Supplementary Tables 7-8) did not suggest much spatial variation for the relationship between cognitive decline and thickness decline, and no obvious systematic variation in this relationship for area.

From a biological perspective, the observed dissociation between thickness and area is consistent with evidence that these measures reflect partly distinct neurobiological processes. Thickness has been linked to neuronal growth and dendritic structures, reduction of synaptic spines and the length of myelinated axons – features known to change with ageing (Fjell and Walhovd 2010; Vidal-Pineiro et al. 2020). Age-related reductions in thickness may reflect cumulative microstructural alterations associated with reduced information processing efficiency. Although causal inference should be made with care, the temporal ordering between baseline thickness and subsequent cognitive change is at least compatible with such a causal interpretation. In contrast, while area does clearly decline with age, the association of this decline with cognitive decline is weaker and less consistent than that for thickness. This pattern also aligns with neurobiological basis of cortical area, which is largely determined by early developmental mechanisms, including progenitor cell proliferation and cortical folding area, consistent with its mediation of polygenic scores on cognition. The formal test of dissociation was not significant, but the pattern is consistent with the view that area, while declining with age, may index earlier genetically influenced cognitive architecture, such that baseline adult levels relate to baseline cognition, whereas subsequent age-related reductions may occur largely independently of cognitive decline. Although we have examined the relationship between cognition and thickness versus area in adults (particularly middle- to old-aged adults in HABS-HD and OASIS-3), the relationship could differ during development (e.g., 0-18 years of age).

Postmortem studies suggest that healthy aging is not necessarily characterised by substantial neuronal loss. Instead, age-related changes are often observed in dendritic architecture and synaptic spine density, with effects varying across cortical layers, regions and individuals (Petanjek et al. 2008; Petanjek et al. 2011). This cellular-level heterogeneity may contribute to the modest associations observed between cortical thickness and cognition across cohorts. These findings are consistent with the view that MRI-derived cortical thinning reflects the cumulative influence of multiple microstructural processes rather than a simple loss of neuronal tissue.

Evidence from neurodegenerative and neuropsychiatric disorders also suggests that cognitive dysfunction can arise from selective alterations in specific neuronal populations and cortical microcircuits, rather than from uniform neuronal loss. In Alzheimer’s disease, particular subsets of corticocortical pyramidal neurons – especially large neurons in layers III and V of association cortices – appear especially vulnerable to degeneration, whereas other neuronal populations are relatively preserved (Hof et al. 1990). Similarly, studies of schizophrenia have reported layer-and cell-type-specific abnormalities, including reductions in the somal size and connectivity of deep layer III pyramidal neurons in the prefrontal cortex, without evidence for widespread cortical neuron loss (Pierri et al. 2001). Together, these findings indicate that morphological measures such as cortical thickness provide only a partial index of the neural alterations relevant for cognitive function, as comparable cortical measurements may arise from distinct underlying cellular and circuit-level changes.

In terms of other caveats, it is important to note that we only focused on cortical measures, and age-related cognitive change is likely to be influenced by the changes in subcortical structures, e.g. hippocampus or other subcortical regions, contributing to variance not captured by thickness or surface area alone. We also assumed a linear dependency between brain change and cognition change, but recent evidence suggests that the association between brain atrophy and memory decline strengthens with age, and primarily affects individuals with above-average brain structural decline (Vidal-Piñeiro et al. 2025), which might be the case for *g* as well (Griffin et al. 2026). Furthermore, polygenic scores were only available in Cam-CAN, preventing replication of the genetic dissociation, and it should be noted that there is debate on the extent of genetic independence of thickness and area (Hofer et al. 2020; van der Meer et al. 2020). Finally, while we found no evidence of attrition bias in our longitudinal analyses, we could only do so in HABS-HD cohort due to either the number or structure of waves.

In summary, the present findings support the claim that thickness and area contribute differently to cognitive ageing, with thickness showing stronger age-related decline, tighter coupling to cognitive change, and greater predictive value for future cognition. Distinguishing these morphometric components clarifies inconsistencies in prior findings and underscores the importance of modelling thickness and area separately when characterising individual trajectories of cognitive ageing.

## Data availability

R code (for all analyses) and data (for Cam-CAN only) are available here: https://github.com/InaDemetriou/GrayMatter_THK_SA_Dissociation

Raw data for OASIS-3 and HABS-HD can be requested from: https://www.nitrc.org/projects/oasis3 and https://ida.loni.usc.edu/home/projectPage.jsp?project=HABS_HD respectively.

## Authors’ contributions

I.D.: Conceptualisation, Methodology, Software, Validation, Formal analysis, Data curation, Visualisation, Writing – Original Draft, Writing – Review & Editing; M.C.: Software, Data curation, Writing – Review & Editing; D.V.P.: Data curation, Writing – Review & Editing; E.E: Data curation, Writing – Review & Editing; S.E.F.: Data curation, Writing – Review & Editing; V.E.P.: Data curation, Writing – Review & Editing; M.B.S.: Data curation, Writing – Review & Editing; D.A.: Data curation, Writing – Review & Editing; A.A.: Data curation, Writing – Review & Editing; T.E.: Data curation, Writing – Review & Editing; R.H.: Conceptualisation, Methodology, Software, Validation, Formal analysis, Resources, Data curation, Writing – Review & Editing, Supervision, Project administration, Funding acquisition.

## Acknowledgements

We thank the Cam-CAN respondents and their primary care teams in Cambridge for their participation in this study, and colleagues at the MRC Cognition and Brain Sciences Unit MEG and MRI facilities for their assistance.

We thank the Cam-CAN team: Project principal personnel: Richard N Henson, Lorraine K Tyler, Kamen A Tsvetanov, Carol Brayne, Edward T Bullmore, Andrew C Calder, Rhodri Cusack, Tim Dalgleish, John Duncan, Fiona E Matthews, William D Marslen-Wilson, James B Rowe, Meredith A Shafto, Marta Correia; Research Associates: Karen Campbell, Teresa Cheung, Simon Davis, Linda Geerligs, Rogier Kievit, Anna McCarrey, Abdur Mustafa, Darren Price, David Samu, Jason R Taylor, Matthias Treder, Janna van Belle, Nitin Williams, Daniel Mitchell, Ethan Knights, Adam Attaheri, Dace Apsvalka, Maite Crespo-Garcia; Research Assistants: Lauren Bates, Tina Emery, Sharon Erzinçlioğlu, Andrew Gadie, Sofia Gerbase, Stanimira Georgieva, Claire Hanley, Beth Parkin, David Troy, Ina Demetriou, Will Duckett; Affiliated Personnel: Matt Bracher-Smith, Valentina Escott-Price, Tibor Auer, Lu Gao, Emma Green, Rafael Henriques; Research Interviewers: Jodie Allen, Gillian Amery, Liana Amunts, Anne Barcroft, Amanda Castle, Cheryl Dias, Jonathan Dowrick, Melissa Fair, Hayley Fisher, Anna Goulding, Adarsh Grewal, Geoff Hale, Andrew Hilton, Frances Johnson, Patricia Johnston, Thea Kavanagh-Williamson, Magdalena Kwasniewska, Alison McMinn, Kim Norman, Jessica Penrose, Fiona Roby, Diane Rowland, John Sargeant, Maggie Squire, Beth Stevens, Aldabra Stoddart, Cheryl Stone, Tracy Thompson, Ozlem Yazlik; and administrative staff: Dan Barnes, Marie Dixon, Jaya Hillman, Joanne Mitchell, Laura Villis.

We thank The Health and Aging Brain Study (HABS-HD) Study Team. The content is solely the responsibility of the authors and does not necessarily represent the official views of the National Institutes of Health. HABS-HD MPIs: Sid E O’Bryant, Kristine Yaffe, Arthur Toga, Robert Rissman, & Leigh Johnson; and the HABS-HD Investigators: Meredith Braskie, Kevin King, James R Hall, Melissa Petersen, Raymond Palmer, Robert Barber, Yonggang Shi, Fan Zhang, Rajesh Nandy, Roderick McColl, David Mason, Bradley Christian, Nicole Phillips, Stephanie Large, Joe Lee, Badri Vardarajan, Monica Rivera Mindt, Amrita Cheema, Lisa Barnes, Mark Mapstone, Annie Cohen, Amy Kind, Ozioma Okonkwo, Raul Vintimilla, Zhengyang Zhou, Michael Donohue, Rema Raman, Matthew Borzage, Michelle Mielke, Beau Ances, Ganesh Babulal, Jorge Llibre-Guerra, Carl Hill and Rocky Vig.

We thank Rogier A. Kievit for the statistical advice and his valuable insights in shaping the analyses. We thank Lihua Xia for providing processed diagnostic labels for HABS-HD.

For the purpose of open access, the author has applied a Creative Commons Attribution (CC BY) licence to any Author Accepted Manuscript version arising from this submission.

## Funding

Cam-CAN was initially supported by the Biotechnology and Biological Sciences Research Council Grant BB/H008217/1, then received further contributions from the European Union’s Horizon 2020 research and innovation programme (‘LifeBrain’, Grant Agreement No. 732592) and intramural funding to the MRC Cognition & Brain sciences Unit [SUAG/046/G101400].

Data were provided in part by OASIS-3: Longitudinal Multimodal Neuroimaging: Principal Investigators: T. Benzinger, D. Marcus, J. Morris; NIH P30 AG066444, P50 AG00561, P30 NS09857781, P01 AG026276, P01 AG003991, R01 AG043434, UL1 TR000448, R01 EB009352.

Data were provided in part by HABS-HD, supported by the National Institute on Aging of the National Institutes of Health under Award Numbers R01AG054073, R01AG058533, R01AG070862, P41EB015922 and U19AG078109.

