## Supplementary Materials for "Dissociable contributions of cortical thickness and surface area to cognitive ageing: evidence from multiple longitudinal cohorts"

*Supplementary Table 1.* Indirect effects of age and polygenic score on cognition through cortical thickness and surface area across cohorts with YoE.

| Cohort | N | Path | Mediator | β | p | Effect % |
| --- | --- | --- | --- | --- | --- | --- |
| CamCAN | 621 | Age → *g* | Thickness | 0.07 | .004 | 12% |
|  |  |  | Area | 0.04 | .001 | 7% |
|  |  | PGS → *g* | Thickness | 0.01 | .094 | 5% |
|  |  |  | Area | 0.01 | .010 | 11% |
|  |  | YoE → *g* | NA | 0.28 | <.001 | NA |
| OASIS-3 | 981 | Age → *g* | Thickness | 0.14 | <.001 | 28% |
|  |  |  | Area | 0.06 | <.001 | 12% |
|  |  | YoE → *g* | NA | 0.24 | <.001 | NA |
| HABS-HD | 2749 | Age → *g* | Thickness | 0.02 | <.001 | 8% |
|  |  |  | Area | 0.02 | <.001 | 7% |
|  |  | YoE → *g* | NA | 0.61 | <.001 | NA |

Supplementary Table 2. Mediation of age on cognition through cortical thickness and surface area after adding paths from ICV to thickness and area. Note that ICV was approximated by eTIV in the Cam-CAN and HABS-HD cohorts, since segmented ICV was not available.

|  |  | Predicted by | Estimate (% mediated), p-value |
| --- | --- | --- | --- |
| Cohort | Mediator | ~ ICV | Age → g |
| Cam-CAN  (N = 621) | Thickness | β=0.05, p=.217 | β=0.12 (17%), p<.001 |
|  | Area | β=0.78, p<.001 | β=0.05 (7%), p<.001 |
| OASIS-3  (N = 981) | Thickness | β=-0.09, p=.009 | β=0.15 (28%), p<.001 |
|  | Area | β=0.75, p<.001 | β=0.07 (14%), p<.001 |
| HABS-HD  (N = 2749) | Thickness | β=0.12, p<.001 | β=0.04 (18%), p<.001 |
|  | Area | β=0.83, p<.001 | β=0.05 (22%), p<.001 |

*Supplementary Table 3.* Correlated change results from simple models, with (‘Preserved’) and without (‘Adjusted (GAMM)’) adjustments for non-linear age effects using a GAMM for thickness, area and volume.

|  | Non-linear age effects | Thickness change | Area change | Volume change | Steiger’s test  (Thickness vs. Area) |
| --- | --- | --- | --- | --- | --- |
| CamCAN  (n=131) | Preserved | 0.17 (3.1%), p=.046 | 0.01 (<1%), p=.936 | 0.11 (1.2%), p=.220 | z=1.34, p=.181 |
|  | Adjusted (GAMM) | 0.11 (1.3%), p=.195 | -0.05 (<1%), p=.582 | 0.04 (<1%), p = .642 | z=1.24, p=.213 |
| OASIS-3  (n=511) | Preserved | 0.18 (3.1%), p<.001 | 0.04 (<1%), p=.383 | 0.11 (1.1%), p=.017 | z=2.51, p=.012 |
|  | Adjusted (GAMM) | 0.14 (1.9%), p=.002 | -0.01 (<1%), p=.742 | 0.09 (<1%), p=.046 | z=2.70, p=.007 |
| HABS-HD  (n=1252) | Preserved | 0.10 (1.0%), p<.001 | 0.08 (<1%), p=.008 | 0.11 (1.2%), p<.001 | z=0.65, p=.515 |
|  | Adjusted (GAMM) | 0.10 (1.0%), p<.001 | 0.06 (<1%), p = .036 | 0.11 (1.1%), p<.001 | z=1.13, p=.260 |

*Supplementary Table 4.* Associations between cognitive change and cortical change/baseline from extended longitudinal models, estimated with and without adjustment for non-linear age effects. Linear Hypothesis Test (LHT) results indicate whether change–change associations differ significantly between cortical thickness and surface area.

|  | N | Non-linear  age effects | Thickness change | Thickness baseline | Area  change | Area  baseline | LHT |
| --- | --- | --- | --- | --- | --- | --- | --- |
| CamCAN | 131 | Preserved | 0.18 (3.4%), p=.037 | 0.24 (6%), p=.005 | 0.004 (<1%), p=.964 | 0.08 (<1%) p=.335 | F=2.09, p=.150 |
|  |  | Adjusted (GAMM) | 0.12 (1.5%), p=.167 | 0.10 (<1%), p=.282 | -0.03 (<1%), p=.701 | 0.04 (<1%), p=.682 | F=1.70, p=.195 |
| OASIS-3 | 511 | Preserved | 0.18 (3.4%), p<.001 | 0.23 (5.5%), p<.001 | -0.01 (<1%), p=.766 | -0.02 (<1%), p=.576 | F=8.33, p=.004 |
|  |  | Adjusted (GAMM) | 0.16 (2.5%), p<.001 | 0.17 (2.9%), p<.001 | -0.05 (<1%), p=.260 | -0.05 (<1%), p=.268 | F=9.52, p=.002 |
| HABS-HD | 1252 | Preserved | 0.12 (1.3%),  p <.001 | 0.12 (1.4%),  p<.001 | 0.05 (<1%),  p=.061 | -0.004 (<1%),  p=.889 | F=2.21,  p=.137 |
|  |  | Adjusted (GAMM) | 0.12 (1.3%), p<.001 | 0.11 (1.1%), p<.001 | 0.04 (<1%), p=.151 | -0.001 (<1%), p=.973 | F = 3.29, p = .070 |

Supplementary Table 5. Bivariate latent change score model results for latent change covariance and cross-lagged relationships between baseline and change measure without (top) and with (bottom) adjustment for baseline age (dg ~ A0 & dThK or dArea ~ A0).

| Cohort | N | Measure | dCortex–dg | Cortex–dg | g–dCortex | dCortex–A0 | dg–A0 |
| --- | --- | --- | --- | --- | --- | --- | --- |
| CamCAN | 131 | Thickness | 0.13, p=.149 | 0.04, p=.662 | -0.02, p=.806 | -0.31, p=.003 | -0.45, p<.001 |
|  |  | Area | -0.04, p=.720 | 0.09, p=.131 | 0.07, p=.447 | -0.05, p=.612 | -0.46, p<.001 |
| OASIS-3 | 137 | Thickness | 0.25, p=.003 | 0.20, p=.020 | 0.15, p=.053 | -0.22, p=.018 | 0.06, p=.474 |
|  |  | Area | -0.01, p=.909 | 0.01, p=.889 | 0.11, p=.223 | -0.29, p<.001 | -0.04, p=.624 |
| HABS-HD | 553 | Thickness | 0.17, p<.001 | 0.05, p=.331 | 0.09, p=.016 | -0.14, p<.001 | -0.19, p<.001 |
|  |  | Area | 0.12, p=.008 | -0.02, p=.711 | 0.12, p=.005 | -0.12, p=.002 | -0.20, p<.001 |

*Supplementary Table 6.* Latent growth model* results for latent change covariance and cross-lagged relationships between baseline and change. sCortex – latent slope for morphometric feature, iCortex – intercept (baseline) for morphometric feature; sg – latent slope for cognition, ig – intercept for cognition.

| Cohort | N | Measure | sCortex–sg | iCortex–sg | ig–sCortex |
| --- | --- | --- | --- | --- | --- |
| OASIS-3 | 137 | Thickness | 0.25, p=.013 | 0.21, p=.012 | 0.29, p=.001 |
|  |  | Area | -0.01, p=.944 | 0.04, p=.673 | 0.39, p=.038 |
| HABS-HD | 553 | Thickness | 0.21, p<.001 | 0.13, p=.005 | 0.10, p=.024 |
|  |  | Area | 0.25, p=.002 | -0.05, p=.232 | 0.21, p=.003 |

* The variances of observed measures were constrained to a small value (0.01) to correct for occasional negative variances (so-called “Heywood cases”, a problem and solution both common in latent growth modelling of behavioural constructs).

*Supplementary Figure 1.* Visual representation of latent growth models in the OASIS-3 and HABS-HD cohorts. Circles denote latent variables: baseline (intercept) levels (i g; i ThK (thickness) / i Area) and latent change factors, i.e. ‘slopes’ (s g; s ThK / s Area). Rectangles represent observed indicators of g at selected three time points (g1, g2, g3) and cortical metrics (Thk1-Thk3; Area1-Area3). Straight single-headed arrows indicate regression paths; curved double-headed arrows indicate covariances. Dashed paths represent parameters fixed for model identification. Coloured paths show the cross-domain coupling effects of primary interest (orange: thickness models; turquoise: area models). Values are standardised parameter estimates. Statistical significance is indicated as: p < .05 (*), p < .01 (**), p < .001 (***).

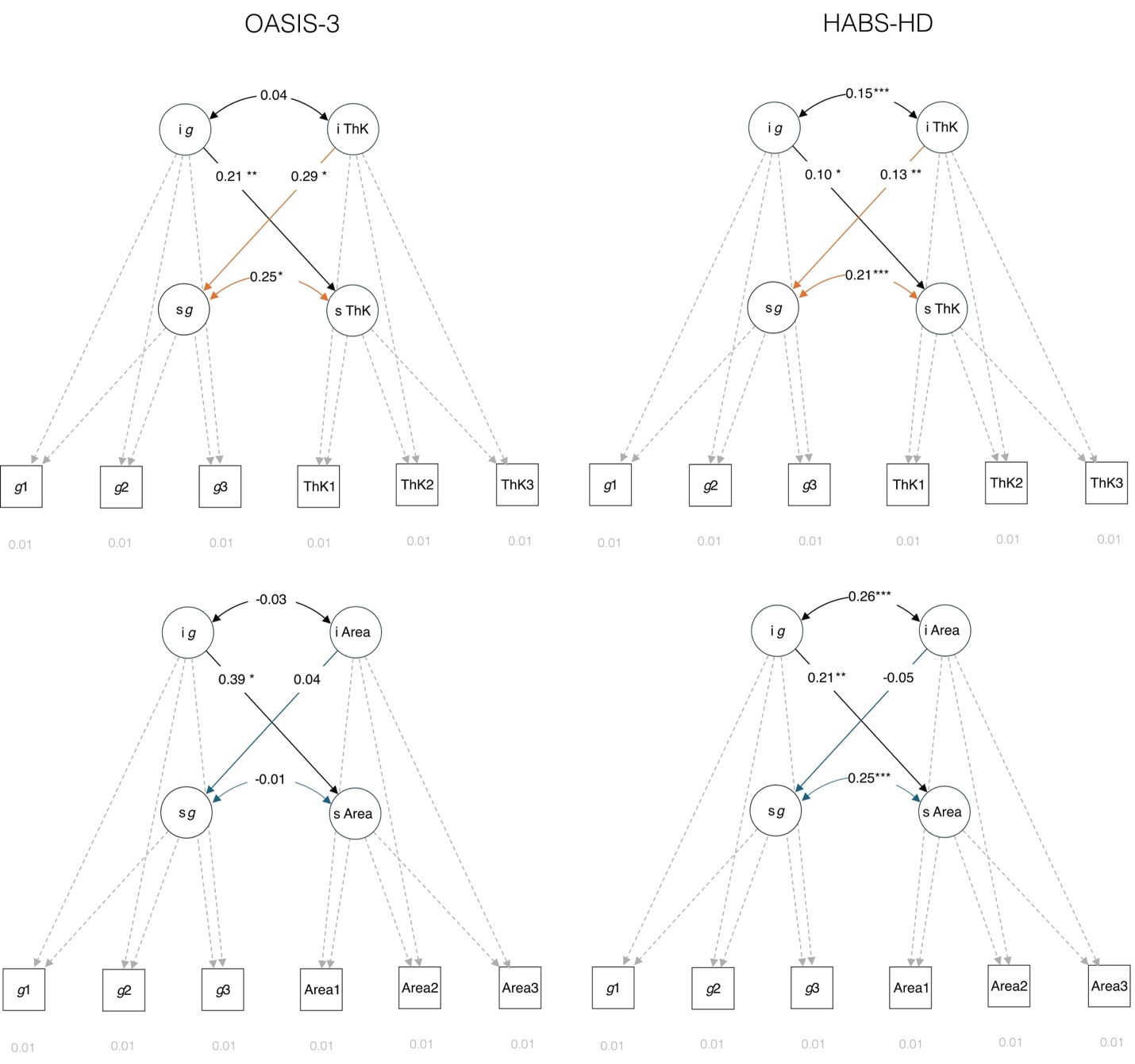

Supplementary Table 7. Region of Interest (ROI) associations between cortical thickness or surface area change and cognitive change across 34 Desikan–Killiany cortical regions (collapsed across hemisphere). P-values are uncorrected; ROIs that survived FDR correction are in bold. ROIs highlighted in grey (first three rows) were those tested by Nyberg et al. (2023). Note that regional area measures were not available in the OASIS-3 cohort.

| Cohort | CamCAN (n=131) | | | | OASIS-3 (n=511) | | | HABS-HD (n= 1252) | | | |
| --- | --- | --- | --- | --- | --- | --- | --- | --- | --- | --- | --- |
| Measure | Thickness | | Area | | Thickness | | Area | Thickness | | Area | |
| ROI | β | p | β | p | β | p | β, p | β | p | β | p |
| inferiorparietal | 0.11 | .203 | 0.08 | .391 | 0.09 | .036 | NA | **0.08** | **.005** | 0.04 | .157 |
| middletemporal | 0.07 | .399 | 0.15 | .077 | **0.17** | **<.001** | NA | 0.06 | .023 | 0.07 | .014 |
| rostralmiddlefrontal | 0.04 | .620 | -0.05 | .537 | 0.06 | .195 | NA | 0.01 | .729 | 0.06 | .023 |
| bankssts | 0.03 | .693 | <.001 | .969 | 0.07 | .099 | NA | 0.06 | .030 | 0.04 | .180 |
| caudalanteriorcingulate | -0.03 | .717 | 0.10 | .268 | 0.02 | .610 | NA | 0.02 | .466 | 0.04 | .207 |
| caudalmiddlefrontal | 0.07 | .445 | -0.10 | .238 | 0.00 | .997 | NA | 0.06 | .048 | 0.07 | .011 |
| cuneus | 0.17 | .052 | 0.05 | .553 | 0.09 | .046 | NA | **0.07** | **.013** | 0.02 | .487 |
| entorhinal | 0.21 | .014 | -0.16 | .074 | **0.18** | **<.001** | NA | **0.07** | **.013** | 0.04 | .199 |
| frontalpole | -0.01 | .911 | 0.11 | .200 | 0.06 | .182 | NA | 0.01 | .668 | 0.03 | .219 |
| fusiform | 0.14 | .102 | 0.08 | .346 | 0.08 | .076 | NA | **0.11** | **<.001** | -0.02 | .415 |
| inferiortemporal | 0.11 | .229 | 0.05 | .559 | 0.11 | .012 | NA | **0.09** | **<.001** | -0.01 | .621 |
| insula | 0.06 | .500 | 0.11 | .219 | 0.09 | .035 | NA | **0.11** | **<.001** | 0.03 | .316 |
| isthmuscingulate | -0.03 | .721 | 0.12 | .165 | **0.16** | **<.001** | NA | 0.03 | .287 | 0.05 | .089 |
| lateraloccipital | 0.23 | .009 | 0.01 | .942 | 0.10 | .032 | NA | 0.06 | .040 | 0.02 | .528 |
| lateralorbitofrontal | 0.05 | .557 | 0.04 | .652 | 0.05 | .266 | NA | <.001 | .910 | 0.07 | .013 |
| lingual | 0.22 | .011 | 0.01 | .912 | 0.07 | .102 | NA | 0.03 | .300 | 0.06 | .031 |
| medialorbitofrontal | 0.04 | .626 | 0.14 | .113 | 0.05 | .289 | NA | -0.01 | .852 | 0.04 | .159 |
| paracentral | 0.14 | .102 | -0.17 | .051 | 0.00 | .912 | NA | **0.07** | **.010** | -0.01 | .807 |
| parahippocampal | 0.17 | .055 | 0.11 | .221 | **0.19** | **<.001** | NA | **0.12** | **<.001** | 0.02 | .381 |
| parsopercularis | 0.02 | .783 | -0.03 | .729 | 0.03 | .499 | NA | 0.03 | .327 | 0.07 | .020 |
| parsorbitalis | -0.11 | .194 | 0.20 | .024 | 0.00 | .943 | NA | 0.02 | .442 | 0.02 | .457 |
| parstriangularis | 0.10 | .279 | 0.05 | .581 | 0.04 | .349 | NA | <.001 | .977 | 0.08 | .004 |
| pericalcarine | 0.21 | .016 | -0.01 | .928 | 0.05 | .304 | NA | 0.04 | .161 | 0.05 | .052 |
| postcentral | 0.20 | .025 | -0.18 | .037 | 0.01 | .752 | NA | 0.06 | .026 | <.001 | .883 |
| posteriorcingulate | -0.07 | .423 | 0.15 | .087 | 0.11 | .013 | NA | 0.04 | .122 | 0.05 | .059 |
| precentral | 0.14 | .112 | -0.07 | .439 | 0.04 | .368 | NA | **0.09** | **<.001** | 0.02 | .479 |
| precuneus | 0.19 | .032 | 0.02 | .845 | 0.08 | .084 | NA | **0.08** | **.003** | 0.07 | .010 |
| rostralanteriorcingulate | -0.06 | .530 | 0.04 | .654 | 0.09 | .037 | NA | 0.04 | .166 | 0.03 | .232 |
| superiorfrontal | 0.06 | .525 | -0.13 | .136 | 0.05 | .290 | NA | **0.08** | **.008** | 0.06 | .027 |
| superiorparietal | 0.19 | .034 | <.001 | .973 | 0.08 | .079 | NA | **0.08** | **.005** | 0.03 | .263 |
| superiortemporal | 0.17 | .049 | -0.01 | .941 | **0.14** | **.002** | NA | **0.09** | **.002** | 0.05 | .103 |
| supramarginal | 0.15 | .093 | -0.07 | .444 | 0.08 | .085 | NA | 0.06 | .039 | 0.06 | .044 |
| temporalpole | 0.17 | .054 | 0.11 | .204 | 0.11 | .015 | NA | **0.09** | **.001** | 0.03 | .307 |
| transversetemporal | 0.19 | .031 | 0.06 | .529 | 0.07 | .113 | NA | 0.02 | .421 | <.001 | .989 |

| Cohort | CamCAN (n=131) | | | | OASIS-3 (n=511) | | | HABS-HD (n= 1252) | | | |
| --- | --- | --- | --- | --- | --- | --- | --- | --- | --- | --- | --- |
| Measure | Thickness | | Area | | Thickness | | Area | Thickness | | Area | |
| ROI | β | p | β | p | β | p | β, p | β | p | β | p |
| inferiorparietal | -0.07 | .610 | 0.17 | .221 | -0.04 | .521 | NA | <.001 | .993 | -0.03 | .448 |
| middletemporal | -0.09 | .477 | 0.25 | .028 | 0.10 | .120 | NA | -0.01 | .887 | 0.04 | .183 |
| rostralmiddlefrontal | -0.20 | .126 | -0.09 | .403 | <.001 | .959 | NA | -0.08 | .028 | 0.03 | .300 |
| bankssts | -0.12 | .281 | <.001 | .993 | -0.06 | .287 | NA | <.001 | .906 | 0.01 | .653 |
| caudalanteriorcingulate | -0.12 | .201 | 0.10 | .257 | -0.03 | .545 | NA | -0.01 | .860 | 0.01 | .690 |
| caudalmiddlefrontal | -0.17 | .213 | -0.22 | .082 | **-0.20** | **<.001** | NA | -0.01 | .705 | 0.04 | .189 |
| cuneus | 0.11 | .300 | 0.07 | .524 | 0.03 | .465 | NA | 0.02 | .519 | -0.03 | .352 |
| entorhinal | 0.17 | .066 | -0.18 | .051 | 0.12 | .020 | NA | 0.05 | .092 | 0.03 | .291 |
| frontalpole | -0.06 | .518 | 0.13 | .182 | 0.02 | .735 | NA | -0.03 | .358 | 0.01 | .692 |
| fusiform | 0.05 | .669 | 0.10 | .310 | -0.04 | .416 | NA | 0.07 | .027 | -0.05 | .074 |
| inferiortemporal | -0.02 | .869 | 0.07 | .521 | <.001 | .968 | NA | 0.05 | .177 | -0.04 | .157 |
| insula | -0.03 | .729 | 0.12 | .195 | 0.02 | .716 | NA | 0.08 | .015 | <.001 | .982 |
| isthmuscingulate | -0.15 | .122 | 0.12 | .165 | 0.10 | .038 | NA | -0.02 | .494 | 0.03 | .344 |
| lateraloccipital | 0.21 | .094 | <.001 | .970 | <.001 | .991 | NA | -0.03 | .474 | -0.03 | .357 |
| lateralorbitofrontal | -0.06 | .544 | 0.05 | .633 | -0.02 | .601 | NA | -0.08 | .022 | 0.05 | .086 |
| lingual | 0.18 | .084 | 0.01 | .932 | <.001 | .971 | NA | -0.02 | .549 | 0.04 | .232 |
| medialorbitofrontal | -0.08 | .457 | 0.18 | .080 | 0.01 | .829 | NA | -0.07 | .034 | 0.02 | .443 |
| paracentral | 0.07 | .502 | -0.21 | .030 | -0.10 | .041 | NA | 0.02 | .582 | -0.05 | .109 |
| parahippocampal | 0.11 | .241 | 0.12 | .206 | 0.13 | .006 | NA | **0.10** | **.001** | 0.01 | .797 |
| parsopercularis | -0.24 | .067 | -0.06 | .602 | -0.13 | .021 | NA | -0.08 | .044 | 0.03 | .367 |
| parsorbitalis | -0.25 | .010 | 0.24 | .013 | -0.06 | .218 | NA | -0.03 | .337 | -0.01 | .772 |
| parstriangularis | -0.09 | .486 | 0.08 | .513 | -0.03 | .582 | NA | **-0.12** | **.002** | 0.06 | .108 |
| pericalcarine | 0.17 | .082 | -0.01 | .921 | 0.02 | .727 | NA | <.001 | .877 | 0.03 | .420 |
| postcentral | 0.16 | .280 | -0.25 | .013 | -0.10 | .042 | NA | -0.02 | .630 | -0.06 | .074 |
| posteriorcingulate | -0.18 | .057 | 0.19 | .059 | 0.05 | .293 | NA | 0.01 | .836 | 0.02 | .471 |
| precentral | -0.01 | .933 | -0.13 | .278 | -0.10 | .063 | NA | 0.04 | .358 | -0.03 | .341 |
| precuneus | 0.13 | .381 | 0.02 | .850 | -0.08 | .190 | NA | 0.01 | .772 | 0.04 | .334 |
| rostralanteriorcingulate | -0.13 | .150 | 0.05 | .647 | 0.05 | .254 | NA | 0.01 | .816 | 0.01 | .765 |
| superiorfrontal | -0.24 | .103 | -0.26 | .033 | -0.10 | .076 | NA | 0.03 | .374 | 0.03 | .429 |
| superiorparietal | 0.12 | .353 | -0.02 | .901 | -0.05 | .406 | NA | <.001 | .996 | -0.04 | .296 |
| superiortemporal | 0.09 | .558 | -0.03 | .847 | 0.02 | .731 | NA | 0.04 | .329 | <.001 | .943 |
| supramarginal | <.001 | .990 | -0.16 | .225 | -0.08 | .175 | NA | -0.07 | .148 | <.001 | .448 |
| temporalpole | 0.11 | .232 | 0.13 | .184 | 0.02 | .673 | NA | 0.07 | .023 | 0.01 | .183 |
| transversetemporal | 0.14 | .141 | 0.06 | .528 | -0.03 | .585 | NA | -0.03 | .389 | -0.02 | .300 |

Supplementary Table 8. Region of Interest (ROI) associations between cortical thickness or surface area change and cognitive change across 34 Desikan–Killiany cortical regions (collapsed across hemisphere), controlled for global thickness and area respectively. P-values are uncorrected; ROIs that survived FDR correction are in bold. ROIs highlighted in grey (first three rows) were those tested by Nyberg et al. (2023). Note that regional area measures were not available in the OASIS-3 cohort.
